# Blastula stage specification of avian neural crest

**DOI:** 10.1101/705731

**Authors:** Maneeshi S. Prasad, Eileen Uribe-Querol, Jonathan Marquez, Stephanie Vadasz, Nathan Yardley, Patrick B. Shelar, Rebekah M. Charney, Martin I. Garcia-Castro

## Abstract

Cell fate specification defines the earliest steps towards a distinct cell lineage. Neural crest, a multipotent stem cell population, is thought to be specified from the ectoderm, but its varied contributions defy canons of segregation potential and challenges its embryonic origin. Aiming to resolve this conflict, we have assayed the earliest specification of neural crest using blastula stage chick embryos. Specification assays on isolated chick epiblast explants identify an intermediate region specified towards the neural crest cell fate. Furthermore, low density culture suggests that the specification of intermediate cells towards the neural crest lineage is independent of contact mediated induction. Finally, we have validated the regional identity of the intermediate region towards the neural crest cell fate using fate map studies in blastula stage chick embryos. Our results suggest a model of neural crest specification at blastula stage, with restricted ectoderm and mesoderm capacities.

## Introduction

Through the evolution of trilaminar, bilaterian embryos, a unique cell population arose in vertebrates termed the neural crest (NC). These cells contribute to the craniofacial skeleton and peripheral neurons and glia – key features of vertebrate lifestyle, and therefore NC are considered to be at the center of vertebrate evolution and diversity. NC is a transient, early embryonic multipotent stem cell population that defines vertebrates through its contribution to key features for the predatory lifestyle including a larger brain enclosure, jaws, and paired sense organs (Gans and Northcutt, 1983; Glenn Northcutt, 2005; Le Douarin, 1980; Le Douarin and Kalcheim, 1999). Improper NC development leads to a host of pathologies known as neurocrestopathies (Bolande, 1996; 1974; Etchevers et al., 2006; Farlie et al., 2004). NC cells are thought to be derived from the ectoderm, which is consistent with their contribution to skin melanocytes and peripheral neurons and glia. However, their ectomesenchymal contributions in the head region – including bone, cartilage, and fat cells, which in other parts of the body are derived from the mesoderm – have been a topic of scientific focus and discourse. If NC cells are truly ectodermally derived, then their contribution to mesodermal-like derivatives suggests that they uniquely defy current assumptions of sequential segregation and restriction of potential. A break from the conceptual cannons of NC induction came in 2006, with the suggestion that NC specification is ongoing during gastrulation, and that it occurs independent from mesodermal or neural contributions (Basch et al., 2006). Accordingly, a pre-gastrula NC would not be subjected to the expected fate restrictions imposed on the three germ layers. Further support for pre-gastrula specification of the NC emerged from other researchers (Patthey et al., 2008a; 2008b). In *Xenopus*, a specified region of NC was shown to exist coincident with the completion of gastrulation (Mancilla and Mayor, 1996). Furthermore, mammalian work using rabbit embryos and a human model of NC formation based on embryonic stem cells (Betters et al., 2018; Leung et al., 2016), also suggest that early anterior NC is specified prior to gastrulation and independent from definitive neural and mesodermal tissues. In *Xenopus*, a specified region of NC was shown to exist coincident with the completion of gastrulation (Mancilla and Mayor, 1996); however, recent work suggests a pre-gastrula origin of NC, and proposed that prospective NC retain stemness markers and pluripotency from epiblast cells (Buitrago-Delgado et al., 2015). These finding highlight the need to understand the precise origins of neural crest cells.

Here, we report for the first time the earliest known specification of NC in chick blastula embryos. We have identified a restricted intermediate territory of the blastula epiblast as capable of generating NC when cultured in isolation under non-inductive conditions. This territory contains prospective NC (pNC) cells which develop independently from apparent mesoderm or neural contributions. Importantly, low density cell cultures of dissociated epiblast cells suggest that this early specification has been established prior to the culture of the cells, and that it can proceed in a cell autonomous and/or contact independent fashion. Additionally, the *in vivo* contribution of the intermediate epiblast territory to the neural crest lineage was validated using fate mapping. Our results suggest that the most anterior NC in amniotes arise from the pluripotent epiblast, prior and independent to definitive germ layer formation, which is in alignment with the multipotent character of NC.

## Results

### Restricted epiblast region at blastula stage is specified towards neural crest cell fate

To understand the ontogeny of the NC, we sought to investigate the earliest cell fate decisions that segregates the pNC cells from other cell fates. Previously, we reported on a specified population of NC in the chick gastrula (Basch et al., 2006). In this report, we asked whether NC cells are specified in blastula stage avian embryos, the stage preceding gastrulation and the appearance of the germ layers. Here, specification refers to an initiated path of differentiation towards a specific fate. While the specified cell does not initially express known markers of the tested fate (pre-migratory and migratory NC), they are able to do so after continuing with the specified program. The continuation of the specified program relies on permissive conditions, which if compromised, could prevent the originally specified fate.

Specification can be assessed by culturing isolated regions of the epiblast under a neutral, serum free and defined non-inductive environment. Using this assay, we analyzed whether chick epiblast explants from stage XII (Eyal-Giladi Staging (Eyal-Giladi and Kochav, 1976)) are specified towards NC cell fate. Accordingly, blastula embryos were collected and the underlying hypoblast layer removed. A horizontal strip from the equatorial plane ~250μm above the Koller sickle was then dissected and further cut into 12 explants of ~80-100μm^2^ each (Fig. 1A). The explants were cultured in isolation in a collagen gel under non-inductive conditions (Basch et al., 2006) for 25h or 45h (corresponding to approximately stage HH4+ and HH8, respectively) and assessed for cell fate specification. In the chick, the expression of the transcription factor Pax7 has been reported to begin at gastrula stage HH4+, where it is used as a key marker of early NC development (Basch et al., 2006). We observed that specifically intermediate explants, and not those taken from the most lateral or medial regions, display clear features of NC specification. Unlike explants from other regions, intermediate explants expressed nuclear Pax7 after 25h of culture, and acquire migratory characteristics with positive HNK1 epitope staining and maintained Pax7 expression after 45h of culture (n=5, Fig. 1A). To verify that migratory NC at 45h arise from the same Pax7-positive intermediate explants at 25h, we generated a series of explants in which each explant was bisected diagonally, and the resulting half fragments were cultured for either 25 or 45h (Fig. 1B, n=10). Immunostaining on these explants indicate that the intermediate explants positive for Pax7 expression after 25h of culture are the same that display migrating NC cell marker (HNK1+) at 45h. To further confirm the nature of the proposed pNC specified at blastula stages, we assayed for the expression of additional NC markers in intermediate explants. We observed the expression of Sox9, Snai2, Msx1 and Tfap2a in the same intermediate explants (Fig. 1C). Taken together, these results suggest that a specific population of intermediate epiblast cells are specified in the blastula epiblast, prior to gastrulation, towards a NC cell fate.

**Fig. 1:**
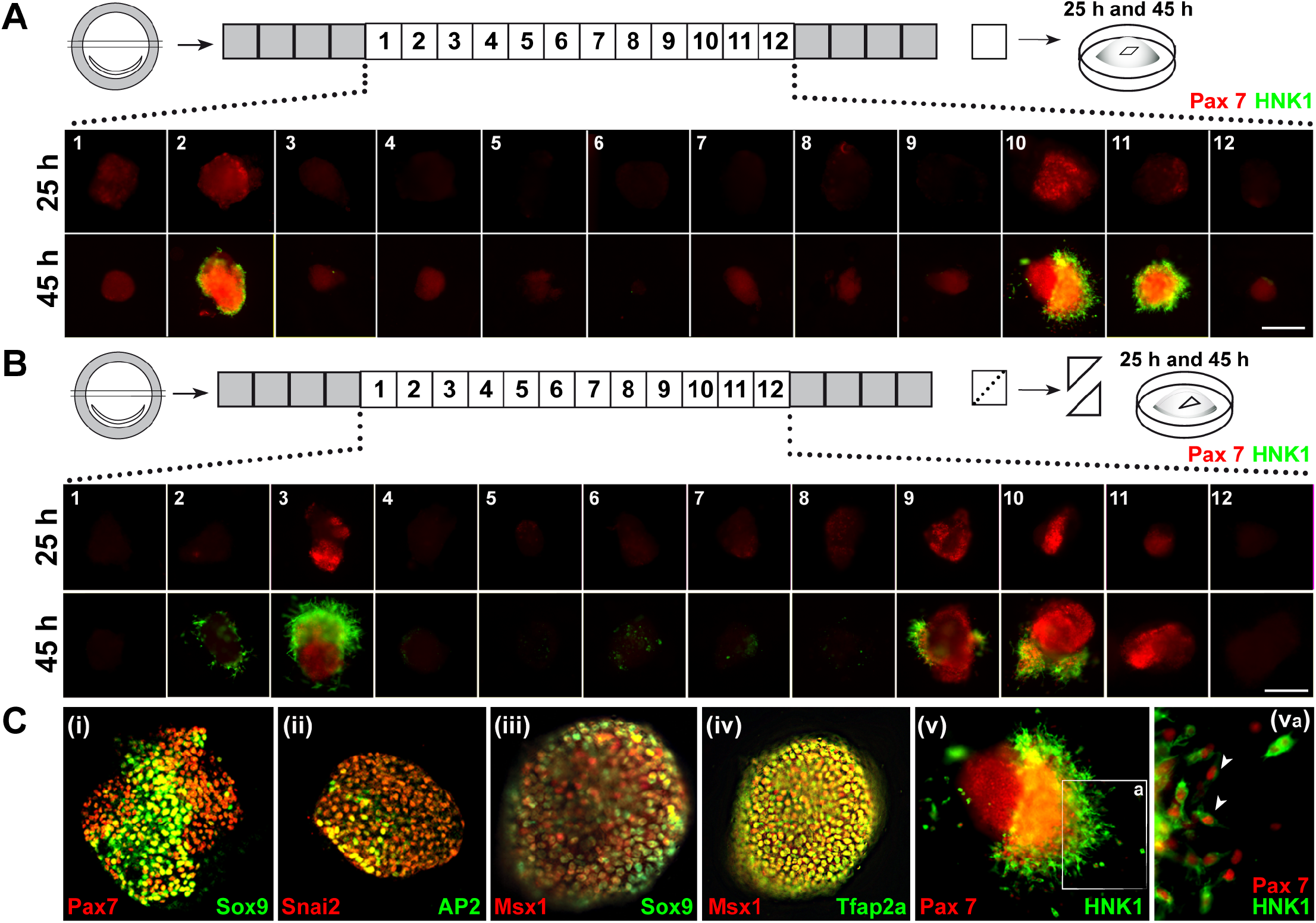
Neural crest cells are specified at blastula stage in a restricted epiblast region. (A) Schematic showing hypoblast-free epiblast explants (≈80μm) from EGK Stage XII chick embryos generated from an equatorial stripe, immersed in collagen gels and cultured under defined non-inducing conditions in isolation. After 25h of culture, lateral explants displayed a robust Pax7+ expression not seen in other explants (n=5). After 45h, a similar intermediate location displays Pax7+/HNK-1+ and clear double positive migratory cells likely to be NCC (n=5; scale bar 100μm). (B) Diagram of a bisection approach to monitor different conditions in original explants. Bisected explants from the same original region generate Pax7+ expressing cells at 25h and migrating Pax7+/HNK-1+ cells at 45h (n=10; scale bar 100μm). (C) Images of intermediate explants with colocalization of neural crest markers at 45h (i) Pax7/Sox9, (ii) Snail2/AP2, (iii) Msx1/2/Sox9, (iv) Msx1/2/AP2, (v) Pax7/HNK-1 explant and (va) magnification of the migrating Pax7+/HNK-1+ cells at 45h.

### Intermediate region of epiblast is specified towards neural crest cell fate independent of neuroectodermal and mesodermal cell fates

Previous work has reported on cell fate specification prior to gastrulation, including neural ectoderm, non-neural ectodermal and mesodermal cell fates (Hatada and Stern, 1994; Onjiko et al., 2015; Patthey et al., 2008b; 2008a; Pegoraro et al., 2015; Shin et al., 2011; Trevers et al., 2018; Wilson et al., 2001). We therefore interrogated the relationship between pNC, neural, and mesodermal tissue in the blastula stage chick embryo. Specification of neuroectodermal (Sox2), mesodermal (TBXT), and NC (Pax7) fates were simultaneously assessed in restricted blastula stage epiblast regions using specification assays described in figure 1A. We observed Sox2 expression predominately in medial explants devoid of robust Pax7 signal (Fig. 2A). While on a few occasions both markers were found in the same explants, with lower levels of Pax7 expression. In the intermediate explants (#2), clear Pax7+/Sox2-expression was identified. Furthermore, only the lateral-most explants displayed TBXT expression, and these explants did not display definitive Pax7 signal (Fig. 2A). Furthermore, the expression of these genes was assessed quantitatively, using RT-qPCR, in the twelve explants after 25hrs of culture. As seen using the immunostaining in figure 2A, intermediate explants were identified with strong Pax7 expression, minimal Sox2, and no TBXT (a lateral mesoderm marker) (Fig. 2B). Together this evidence strongly supports a distinct regional specification towards NC in the intermediate epiblast, with medial and lateral epiblast displaying neural and mesodermal specification, respectively.

**Fig. 2:**
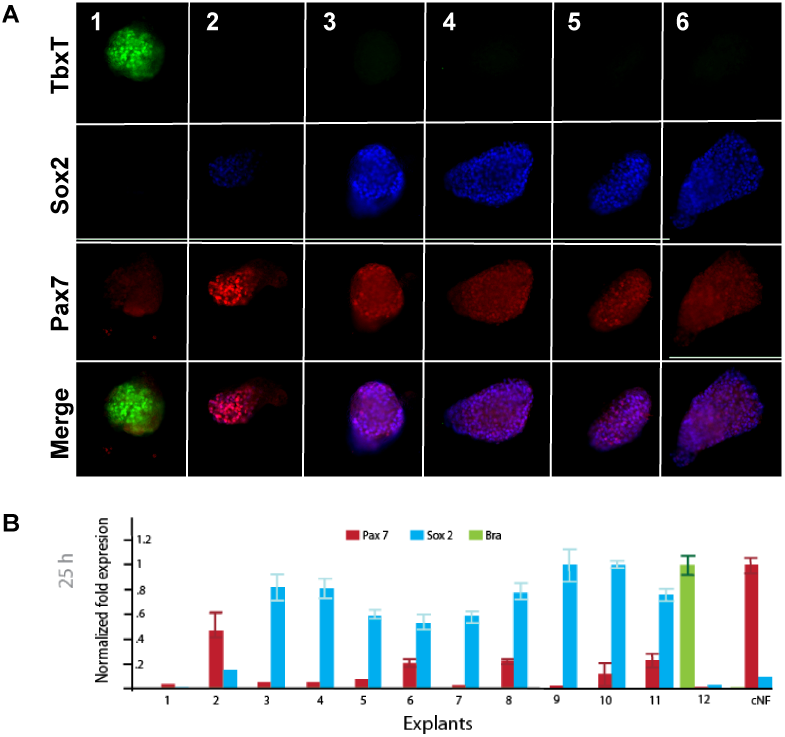
Intermediate region of epiblast is specified towards neural crest cell fate independent of neuroectodermal and mesodermal cell fates. (A) Explants from EGK Stage XII chick embryos, collected after 25h and analyzed for expression of different cell fates; mesodermal (TBXT, (Brachyury) lateral-most), neural (Sox2, medial), and NC (Pax7, intermediate explant). (B) RT-qPCR analysis of twelve explants after 25h of culture, for the expression of Pax7, Sox2, and BraT (TbxT) (a lateral mesoderm marker). Expression is represented as a normalized fold expression to the positive control region for each gene: neural fold (NF) for Pax7 expression and the neural plate for Sox2 expression (not shown). Error bars are standard deviation between technical replicates. Out of the four individual sets of experiments each with 12 explants and controls, two representative RT-qPCR graphs are shown.

### Low density culture of epiblast cells reveals neural crest cell specification at blastula stage

The regional specification of prospective NC in intermediate epiblast explants exposes clear heterogeneity of epiblast cells. This is also clear in the non-homogeneous expression of NC markers in the intermediate explants, and suggests that the intermediate explants are composed of cells of different identities. It seem therefore possible that the identified specification could be the result contact mediated interactions between these cells. To address this possibility, we decided to assess specification in explants subjected to cell dissociation and plated at low density aiming to obtain sufficiently distant cells. The intermediate and medial regions of epiblast from EKG Stage X-XII blastula embryos, were dissociated into single cells using accutase, plated on a layer of collagen at very low density (10-20 cells/ cm^2^) and either fixed immediately (Time 0), or cultured for 30h in neutral media without growth factors (Fig. 3A). These cultures were then evaluated for Pax7 and Sox9 expression via IF. As control, neural fold cells from HH Stage 6 embryo, were dissociated and plated in a similar manner, and processed for IF without culturing them. As expected cells from low density cultures of HH Stage 6 display robust Pax7/Sox9 expression. No Pax7/Sox9 expression was observed in intermediate or medial low density cells at time 0 (right after plating). Instead, nearly 38% of the cells from low cell density cultures from the intermediate epiblast region expressed Pax7, and 24% expressed Sox9 (4 replicates), while cells from medial region contained only 8% Pax7 and 6% Sox9 positive cells (Fig. 3B). The low density dissociation experiment suggests that when cultured in isolation, a certain percentage of intermediate epiblast cells express NC markers in absence of cell-cell contact suggesting a potential cell autonomous specification of neural crest when cultured *in vitro* in absence of *in vivo* repressive cues. This experiment further suggests that a specification program has already been initiated within single cells of the heterogenous blastula epiblast.

**Fig. 3:**
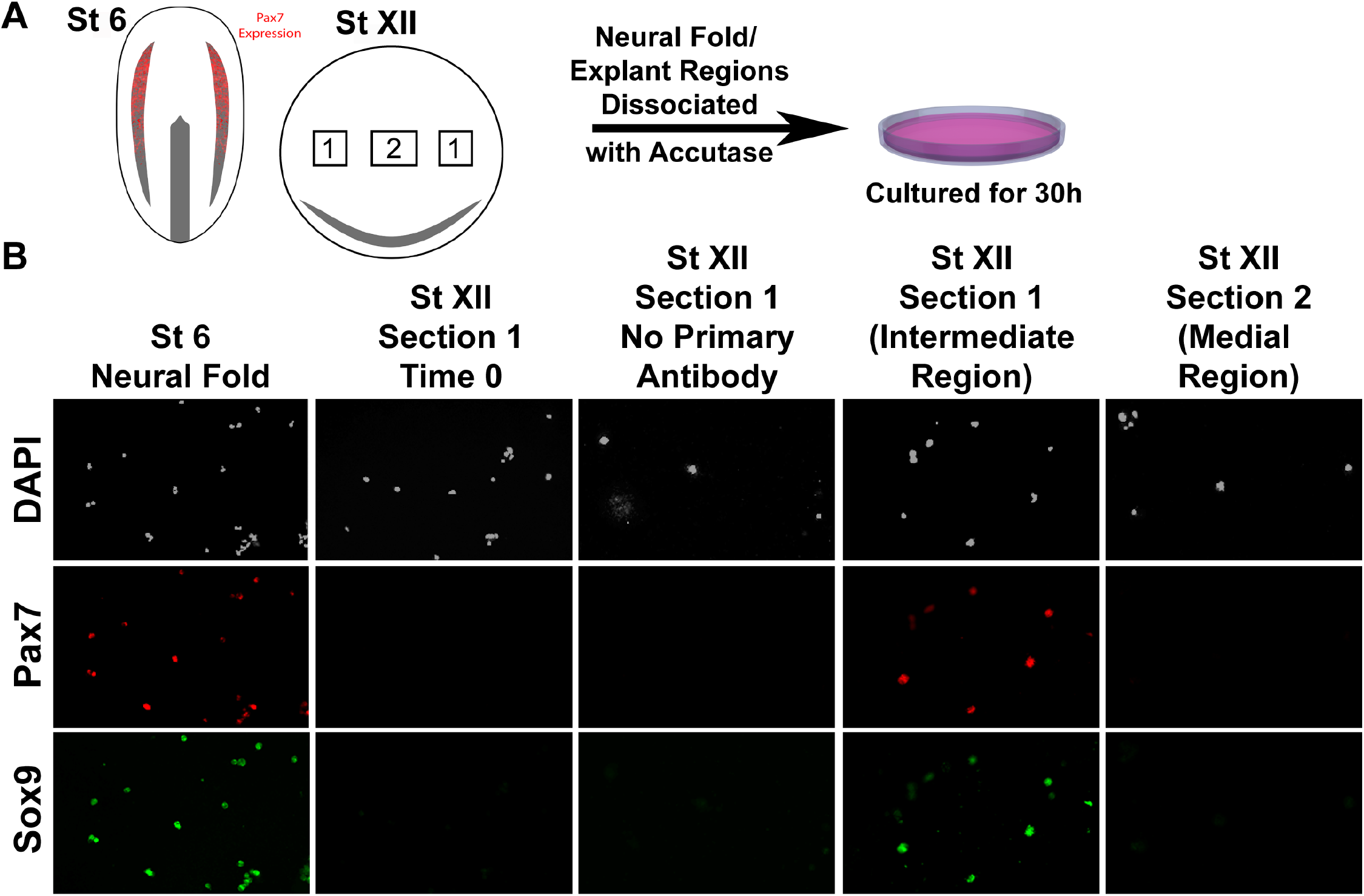
Low density isolated cell analysis of intermediate epiblast region identifies cell autonomous specification of neural crest in culture. (A) Schematic showing HH stage 6 and EGK stage XII embryos used for low density isolated cell analysis and the workflow. Cells were dissociated from Stage 6 embryo marked by Pax7 expressing neural fold region (boxes sections within the red region) (positive control), plated, fixed and immunostained for Pax7/Sox9. Stage XII embryo explants marked as 1 (intermediate) and 2 (medial) regions were dissociated and cultured at very low density (10-20 cells/cm^2^) on a thin layer of collagen gel under non-inducing conditions for 30h. (B) Isolated cells in the low density culture were immunostained for Pax7/Sox9 expression from neural fold region, intermediate region (section 1) at time 0 (immediately after plating), intermediate region (section 1) used as no primary control, and from sections 1 and 2. Intermediate region (section 1) had the highest Pax7 positive cells (38%) with ~8% Pax7+ cells in medial region (n=4).

### Intermediate epiblast cells contribute to neural crest lineage *in vivo*

The explant studies provide a clear and precise approach towards temporal and spatial identification of NC specification. However, to assess the contribution of the specified cells within the explants towards the NC lineage, *in vivo* analysis of the epiblast cells is required. We explored the *in vivo* contribution of the intermediate epiblast cells at stage XII embryos towards NC, using lineage tracing analysis with DiI and DiO, cultured the embryos from 16 to 36h (EC culture) (Chapman et al., 2001) and subsequently fixed and immunostained for Pax7. Labeled regions corresponding to the intermediate explants (Fig. 4B) contributed to the Pax7-expressing neural plate border (NPB) (Fig. 4C’) and neural folds (Fig. 4C”), matching the expected location of NC cells (n= 8/11) (Fig. 4D”). In a few cases, targeted regions labeled simultaneously neural fold/NC along with mesoderm (n=1/11), neural (n=1/11), and non-neural ectoderm (n=3) (summarized in Fig. 4A). In contrast, labeled medial cells contributed to the neural tube (n=3/3) (Fig. 4C’). While mixed contributions were observed from few regions, most of the intermediate epiblast cells populated the NPB and neural folds expressing Pax7, lending *in vivo* support to the notion that at blastula stages the intermediate epiblast harbors pNC cells. Chi-squared test of observed NC contribution from the intermediate region compared to medial region revealed a statistically significant contribution of intermediate epiblast cells towards the NC lineage (p-value=0.024). These results are in agreement with previous lineage tracing studies (Hatada and Stern, 1994) during blastula stage of development. Taken together, our explant specification experiments and lineage tracing studies strongly suggest that cells in a restricted “intermediate” domain of the avian blastula epiblast are already poised to initiate the NC developmental program, well before and thus independently of, definitive mesoderm or neural contributions.

**Fig. 4:**
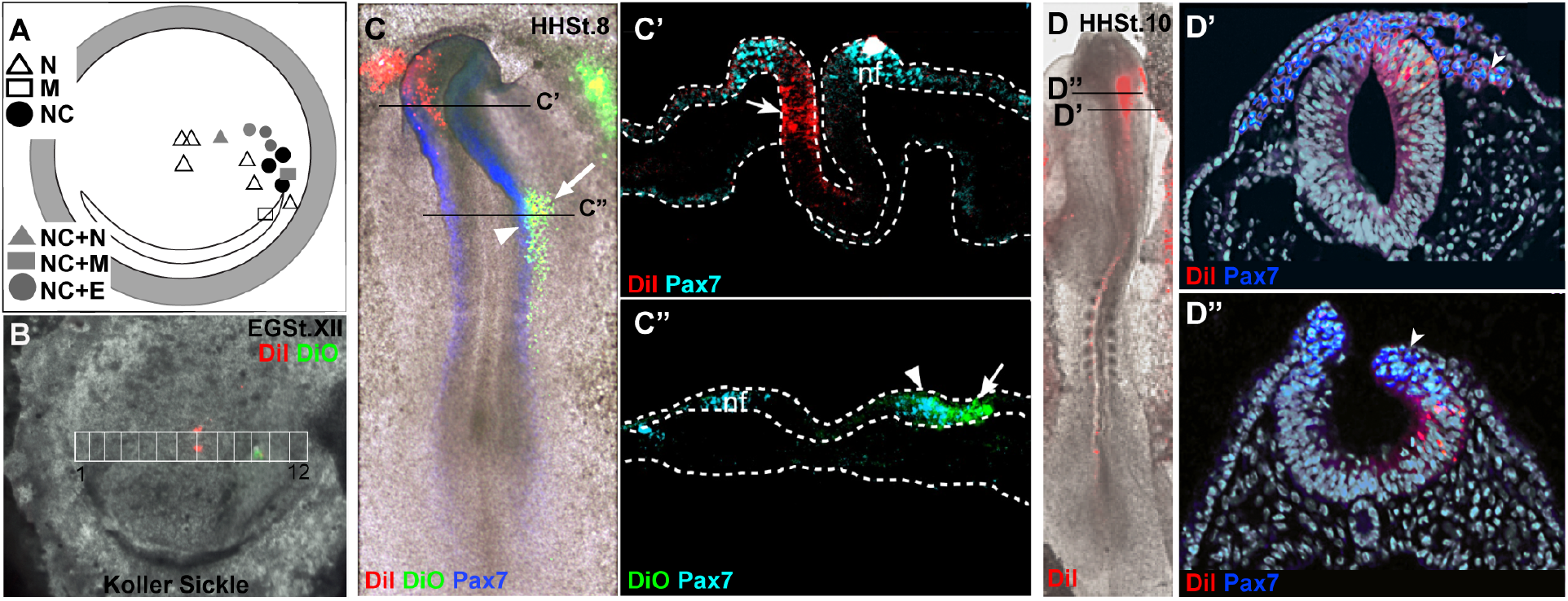
*In vivo* lineage tracing validates the contribution of intermediate epiblast region to neural crest lineage. (A) Schematic summary identifying regions in the epiblast that contributed to NC (n=8) alone (4) or in combination with other fates (N-neural, NC-neural crest, E-epidermal, M-mesodermal). Chi-Squared analysis identified statistically significant (p<0.05) NC contributions of the cells within the intermediate epiblast region. Embryo labeled with DiI/DiO at time 0 (B) and after 28h of culture at St. 8 (C); sections demonstrating contribution to the CNS by cells in the middle of the embryo labeled with DiI (C’, C”) and contribution to the lateral neural fold by cells in the intermediate epiblast labeled with DiO, colocalized with Pax7 expression or lateral to it (arrowhead and arrow respectively, C’, C”). (D) Second embryo labelled with DiI on intermediate region of epiblast, demonstrating anterior neural fold localization of DiI (St. 10). (D’) and (D”) Neural tube cross-section at denoted anterior axial level with colocalization of DiI/Pax7 expression domains in migratory (arrowhead in (D’)) and premigratory (arrowhead in (D”)) neural crest cells and some cells in neural epithelium.

## Discussion

According to the accepted sequential segregation of plasticity, pluripotent cells differentiate and give rise to the three germ layers, endoderm, mesoderm, and ectoderm, each with a distinct potential restricted in comparison to their progenitor. In turn, each of the germ layers differentiate into progenitors with progressively more restricted potential, ultimately generating the specific cell types that constitute the building blocks of the vertebrate body. A recognized exception is the primordial germ cell lineage, which arises independently from gastrulation (Magnusdottir and Surani, 2013; Saitou and Yamaji, 2012). Classic models of NC formation suggest that NC arises from the ectoderm, and therefore, one would expect them to be devoid of mesoderm and endoderm associated potential. However, NC generated ectomesenchymal derivatives encompass ectoderm and mesoderm capacities, and thus represent a difficult paradigm. Efforts to resolve this issue include the suggestion that the NC constitute a fourth germ layer (Hall, 2018; 2000), and a recent model proposing that NC retains stemness markers and the same potential as pluripotent stem cells (Buitrago-Delgado et al., 2015). Our work presented in this report points to a model of blastula stage specification of NC as a segregated population of cells distinct from other cell fates.

We previously showed that anterior cranial neural crest specification is on-going during gastrulation, well before the overt expression of Pax7, the earliest restricted marker associated with NC in chick and rabbit embryos (Basch et al., 2006; Betters et al., 2018). This specification appears to be independent from either mesoderm or neural ectoderm (Basch et al., 2006). In this report, we expose the earliest known specification of anterior cranial NC in the chick blastula embryo. Using epiblast explants at high spatial resolution we demonstrate NC specification in a restricted epiblast region. Furthermore, low density cultures of intermediate epiblast cells suggest NC specification is independent of cell-cell contact mediated inductive interactions in the absence of repressive cues experienced by these cells *in vivo*. However, the role of signaling mediated over long range between the single cells or within the microenvironment of individual single cells cannot be discounted and will require further evaluation during NC specification from epiblast cells. We also observed heterogeneity within the regionalized epiblast, as our single cells dissociation experiments revealed that 66% of the cells within the intermediate explants contributed to Pax7-positive NC cells. The heterogeneity of the early epiblast marked by a progressive loss of pluripotency has previously been documented in multiple species, including chick and human (Chen et al., 2018; De Paepe et al., 2014; Shi et al., 2015). We provide further *in vivo* support for NC specification using fate map studies, and validate that this intermediate epiblast region contains pNC cells that contribute to NC *in vivo*. Our study found contributions from several of our injections to mesoderm, neural ectoderm, and endoderm (with or without co-contribution to NC) which is in agreement with the broad allocation for those fates provided by Hatada and Stern seminal work (Hatada and Stern, 1994). Together, our results from explant, low density single cells and fate mapping very clearly suggest an early specification of prospective NC at blastula stage chick embryos. Importantly, we observe a distinct regional predisposition of epiblast cells to adopt neural, mesodermal or NC fates. While our results do not rule out the possibility of other ectodermal fates arising from the same region of epiblast, our assay clearly demonstrates the propensity of the blastomeres in the intermediate region to be uniquely specified towards NC cell fate. Such blastula stage specification of various cell fates has been previously suggested. The pre-gastrula stage specification of neuroectoderm (Streit et al., 2000; Wilson et al., 2001) and non-neural ectoderm (Wilson et al., 2001) has been documented in chick embryos. These studies have suggested that medial epiblast territories are specified towards the neural cell fate, while the intermediate epiblast explants are fated towards neural plate border (Wilson et al., 2001) and NC (Trevers et al., 2018). Our work is in agreement with these findings and points towards the blastula stage specification of NC, neuroectoderm, and non-neural ectodermal cell fates in the chick embryo (Fig. 5). However, our model does not disregard the possibility of continued NC specification during later developmental stages that might be required for proper NC formation in the presence of multiple repressive cues from the other cells surrounding prospective NC region *in vivo*. This continuation of the NC program may be dependent upon definitive ectodermal and mesodermal contributions.

**Fig. 5:**
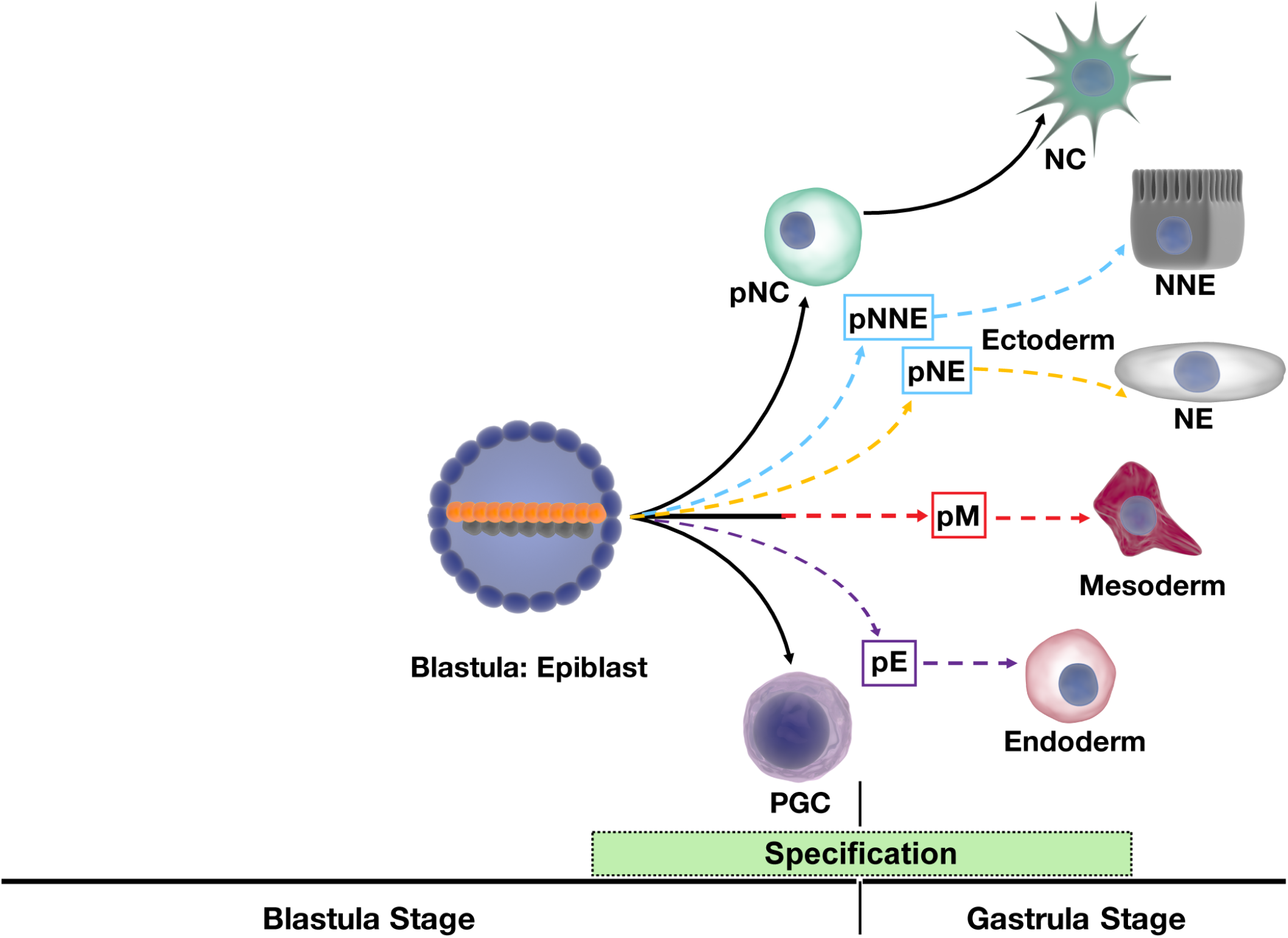
Model of NC cell fate specification from the epiblast. Beginning with early blastula, we propose a model of early NC cell fate specification from epiblast cells (depicted in orange color with underlying hypoblast layer in grey color). Based on our study, the model suggests specification of prospective NC (pNC) from the epiblast during blastula stages of embryonic development. The earlier specification of ectodermal and mesodermal fates suggested by various studies are also reflected in the model. pNE, prospective neural ectoderm; pNNE, prospective non-neural ectoderm; pM, prospective mesoderm; pE, prospective endoderm; PGC, primordial germ cells.

Given the blastula stage specification of NC, the early signaling contributions to specification needs to be addressed in future studies. It has been previously documented that at blastula stage in the lateral/intermediate regions of the epiblast, containing the pNC cells, Wnt/β-catenin is active and required for expression of NPB/NC markers (Wilson et al., 2001), suggesting a potential early role of Wnt/β-catenin signaling during NC specification. Additionally, further analysis of transcriptional changes leading to NC specification from epiblast cells will help in identifying the regulatory network involved during these early steps of NC induction. It is essential to assess this early NC regulatory network, especially under the light of new studies exposing a predisposition of blastomeres towards specific cell fate, based on the relative expression ratios of different lineage specifiers (Shi et al., 2015). To better understand the cell fate specification of different lineages, positional single cell transcriptional analysis at blastula stage is needed to identify the network structure of individual cells to ascertain their predisposed cell fates. Based on our experiments in chick, we propose prospective NC cells as the earliest NC cell state specified during blastula stages from epiblast cells, prior to the formation and concomitant fate segregation associated with the three germ layers. It is believed that chick epiblast and human ES cells are primed, or more advanced than the naïve state (Mak et al., n.d.), and display differentiation bias towards certain cell fates (Mak et al., n.d.; Shin et al., 2011), suggesting an early specification of cell fates prior to gastrulation in multiple species (Sheng, 2015). Our data from chick suggests a similar model for NC specification from epiblast cells prior to gastrulation. This model establishes a direct lineage between epiblast and NC, which are endowed with multipotent potential to form mesectodermal derivatives throughout the vertebrate body plan. Our model provides a parsimonious perspective underscoring the formation of this important vertebrate cell type in amniotes.

## Materials & Methods

### Chicken embryos

Fertile hen eggs were obtained from Hardy’s Hatchery (Massachusetts, USA) and Sunstate Ranch (Sylmar, CA). Embryos were staged according to Hamburger and Hamilton (Hamburger and Hamilton, 1951) for stages 1 and over or Eyal-Giladi and Kochav (EGK) for preprimitive streak (prestreak) stages, Stage IX to XII. Embryos were cultured at 38°C in humidified incubator.

### Chick embryo explant culture

The hypoblast of pre-streak embryos at stage XII was mechanically removed using glass needles, and a horizontal strip of epiblast tissue was cut from the center of the embryo. This strip was trimmed to include only the area pellucida and dissected into 12 equivalent-sized squares (each approximately 100μm^2^) and kept in PB1 buffer (5.97g/L NaCl, 0.2g/L KCl, 1.142g/L NaH2PO4, 0.19 g/L KH2PO4, 0.04 g/L Sodium Pyruvate, 1g/L Glucose, 0.1g/L MgCl2-6 H20, 0.14g/L CaCl2-2H20, 0.06g/L Penicillin, 0.05g/L Streptomycin, 0.01g/L Phenol Red, 4mg/L BSA (added just prior to use)) till next step. These squares were immobilized in separate collagen gels in four-well plates. Collagen gels were prepared by combining 90μl of 3.68 mg/ml collagen (BD Biosciences), 10μl of 10X DMEM (Gibco) and 3.7μl of 7.5% sodium hydrogen carbonate. Collagen gels were immersed in DMEM/F12 containing N2 supplement (Gibco) for 35-48 hours at 37°C. Explants were fixed in 4% paraformaldehyde for 15 minutes before immunostaining.

### Single cell dissociation experiment

The hypoblast of pre-streak embryos at stage XII was mechanically removed using glass needles, and a horizontal strip of epiblast tissue was cut from the center of the embryo. Horizontal epiblast strip was further dissected into 3 sections (2 intermediate and one medial) of ~150cells each (Fig. 2A). The sections were collected in PB1 buffer and treated with accutase for 5min, rinsed in PB1 buffer and dissociated via pipetting. The dissociated cells were washed 2x in PB1 and then transferred in minimal PB1 onto collagen sheets in a chamber slide and was placed in incubator for 15min. Cell density after plating was around 10-20 cells per cm^2^. The chambers were then covered with neutral media (DMEM/F12 + N2 + Pen/Strep + 0.1%BSA) and placed back in the incubator for 30h. The chambers were washed with PBS and fixed in 4% paraformaldehyde for 30min before immunostaining. The number of cells obtained from each section after the complete dissociation and culture procedure were ~20 due to loss during dissociation and plating.

### *In vivo* lineage tracing

At the pre-streak stage, Stage XII, embryos were injected with DiI and DiO (Molecular Probes) into cells of the lateral and medial region of the epiblast layer. Lateral region correspond to the explant #2-3 described above, while medial regions correspond to explant # 6-7. Embryos were cultured at 37°C for 25-48 hours in EC culture (Chapman et al., 2001), then fixed in 4% paraformaldehyde for 15 minutes before immunostaining. Embryos were mounted in gelatin and sectioned at 12μm using a Leica CM1900 Cryostat. Sections were mounted with Permafluor (Thermo Scientific). Images were acquired on Nikon Eclipse 80i microscope, and processed in Adobe Photoshop.

### Immunostaining for chick embryos and explants

Immunostaining for chick embryos and explants were performed as previously described (Basch et al., 2006; Stuhlmiller and Garcia-Castro, 2011). Collagen gels containing explants were fixed with 4% paraformaldehyde for 10 minutes and then washed three times with PBS. Gels were blocked with PBS containing 1% BSA and 0.1%Tween-20 (PBST) for 1 hour at room temperature. Double or triple staining was performed. Primary antibodies for mouse IgG1anti-Pax7 [1:50; Developmental Studies Hybridoma Bank (DSHB)]; mouse IgG1 anti-Msx1,2 (1:50; 4G1, DSHB), mouse IgG1 anti-Snail2 (1:100; 62.1E6, DSHB), mouse IgG2b anti-AP2 (1:50; 3B5, DSHB), mouse IgM anti-HNK-1 (1:100; 1C10, DSHB), goat IgG anti-Sox2 (1:100; R&D AF2018), goat IgG anti-Sox9 (1:100; R&D AF3075) and rabbit IgG anti-Brachyury (1:10) were diluted in PBST and incubated at 4°C overnight. Primary antibody was washed 3 × 10 minutes with PBST. Gels were then incubated with secondary antibodies (goat anti-mouse IgG1Alexa 568, 1:2500; goat anti-mouse IgG2b Alexa 488, 1:2500; donkey anti-IgM Cy5, 1:500; donkey anti-goat IgG 488 or 633, 1:2500 and goat rabbit anti-IgG Alexa 488, 1:2500) diluted in PBST and incubated at 4°C overnight. Secondary antibody was washed 3 × 10 minutes with PT and then stained with DAPI (5μg/mL) for 5 minutes and washed again 3 × 10 minutes with PBS before imaging. Primary antibodies used for chick immunostaining are listed in table 1.

**Table 1:** Antibodies used for immunostaining

| **Antibody Name** | **Vendor** | **Catalog** | **Dilution** | **Species/ Isotype** | **Used in Species** |
| --- | --- | --- | --- | --- | --- |
| MSX1/2 | DSHB | 4G1 | 1:50 | Mouse/IgG1 | Gallus gallus |
| Snai2 | DSHB | 62.1E6 | 1:100 | Mouse/IgG1 | Gallus gallus |
| Tfap2a | DSHB | 3B5 | 1:50 | Mouse/IgG2b | Gallus gallus |
| SOX2 | R&D | AF2018 | 1:100 | Goat/IgG | Gallus gallus |
| SOX9 | R&D | AF3075 | 1:100 | Goat/IgG | Gallus gallus |
| HNK-1 | DSHB | 1C10 | 1:100 | Mouse/IgM | Gallus gallus |
| TbxT<br>(Brachyury) | Gift from Dr Susan Mackem | NA | 1:10 | Rabbit/IgG | Gallus gallus |

### Image processing for chick explants

Images were taken using a Nikon Eclipse 80i microscope and processed in Adobe Photoshop CC version 14.2.1. Images of each experiment were taken in a fluorescent scope (Nikon) using the same settings for each fluorophore channel. Images were compiled in a Photoshop grid image and intensity levels were adjusted at the same time. The threshold levels were set using a positive reference when a clear nuclear staining was detected in an explant of the series.

### Gene expression analysis

#### For chick explant culture

After the treatment, explants were collected in a solution for RNA extraction using the provider specifications (Qiagen RNAeasy). Total RNA was collected in 14μl and was reverse transcribed using the iScript kit (Bio-Rad). Real time was performed in an iQ5 cycler (Bio-Rad) for 40 cycles using SYBRGreen. Three reference genes (β-actin, H2b and H4) were used in each experiment, and Pax7, Sox2, TbxT were the genes analyzed. The data analysis was performed using the ΔΔCT formula, using as control the higher expression. Positive controls of neural plate and neural folds were used to compare the expression of Sox2 and Pax7, respectively.

### Statistical analysis

Chi-Squared test for observed statistical significance for lineage tracing experiments in chick embryo were done using two-way Contingency table, with one-degree of freedom and represented as p-value significance for observed NC contribution of cells labelled within intermediate and medial epiblast regions.

## Acknowledgements

Imaging was performed at the UCR Microscopy and Imaging Core Facility and the UCR Stem Cell Core Facility (California Institute for Regenerative Medicine (CIRM) funded shared facility). We are grateful to Dr. Ken Cho (UC Irvine) and Dr. Ira Blitz (UC Irvine) for helpful conversations, suggestions and critical reading of the manuscript.

## Data and materials availability

All data is available in the main text or the supplementary materials.

